# An Implantable Poly(pro-curcumin) Film for Local, Long-Lasting Neuroprotection After Spinal Cord Injury

**DOI:** 10.64898/2026.01.05.697760

**Authors:** Carlos A. Toro, Nicholas P. Johnson, Zaara Suhail, Ruiwen Chen, John T. Leman, Nikita Waskiewicz, Shahrukh Khalique, Granit Oroshi, Gomathi Jayaraman, Alexey Kozlenkov, Ravi Iyengar, Ryan J. Gilbert, Christopher P. Cardozo, Edmund F. Palermo, Mustafa M. Siddiq

## Abstract

Spinal cord injury (SCI) is a notoriously intractable medical problem for which there is no neuroprotective pharmacotherapy, despite four decades of clinical trials on systemically administered small molecules that target plausible injury mechanisms. The shared failure mode of those trials is delivery: oxidative and inflammatory injury evolves locally at the lesion site over days to weeks, a timescale that far exceeds the half-life of systemically administered drugs. Furthermore, systemically administered drugs often cannot be dosed at high enough levels to achieve efficacious concentrations locally at the lesion site, without off-target effects. In this study, we show the first example of a locally implantable drug depot that exerts long-lasting activity at the lesion site. We demonstrate *in vivo* efficacy of a poly(pro-drug) thin film deployed locally, to address delivery problem in SCI. The films are composed of a copolyester in which curcumin, a potent natural antioxidant, is incorporated as a backbone repeat unit alongside poly(ethylene glycol) sebacate segments. The soft, thin film (∼100 µm) is employed as an epidural implant that quenches reactive oxygen species on-contact and can sustain activity for over two months, which is orders of magnitude longer than free curcumin (half-life of ∼10 min in buffer). A single epidural P50 film, placed on the exposed dura immediately after a moderate-to-severe T9 contusion in mice, produced a statistically significant improvement in Basso Mouse Scale locomotor recovery at 14 days post-injury, relative to a non-antioxidant control polymer of same architecture. The film preserved myelinated white matter, attenuated reactive gliosis (GFAP, Iba1), and produced coherent downregulation of neuroinflammatory gene-expression programs. These effects were spatially restricted to tissue rostral to the lesion, consistent with on-contact, locally confined polymer activity. Because curcumin is covalently enchained within the polymer itself, rather than blended as a payload in a carrier matrix, release is governed by backbone hydrolysis and the architecture can generalize to other phenolic bioactives, potentially establishing epidural poly(pro-drug) thin films as a mechanistically grounded, surgical platform for sustained local pharmacotherapy of CNS injury.

## INTRODUCTION

Spinal cord injury (SCI) is a devastating condition with a complex and prolonged pathophysiology that unfolds over weeks to months following the initial trauma.^1^ More than four decades of pharmacological clinical trials in spinal cord injury (SCI) have failed to yield a single approved neuroprotective therapy. These trials have employed methylprednisolone (NASCIS I– III), GM-1 ganglioside (Sygen), naloxone, tirilazad, riluzole (RISCIS), minocycline, and Cethrin/VX-210, among others. The mechanistic targets in these trials were not unreasonable: oxidative stress, glutamate excitotoxicity, lipid peroxidation, calcium-mediated necroptosis, and Rho-pathway-driven growth cone collapse all participate verifiably in the injury cascade that propagates from the impact site over hours-to-weeks following the initial mechanical insult.^2^ The most plausible reason why these trials all failed is that systemically-administered small molecules with short plasma half-lives, narrow therapeutic windows, and limited blood–spinal-cord-barrier penetration cannot maintain efficacious drug concentrations at the lesion site across the multi-week window during which secondary injury evolves. Furthermore, escalating systemic dose to reach the spinal cord can lead to adverse effects which negate any benefits. In contrast, a delivery modality that places a sustained, local source of neuroprotective drug *directly at the lesion site for extended release* and that can be deployed, using current surgical workflow for decompression and stabilization, has therefore become an important translational requirement for the field. The tissue segment, on which such a delivery system must act, is well characterized. In the acute phase after SCI, microhemorrhages and progressive cord swelling within the rigid spinal canal produce localized ischemia and impaired autoregulation of blood flow. ^3–5^ The ensuing secondary cascade is dominated by release of toxic excitatory neurotransmitters, inflammatory cytokines, and oxidative stress, all of which drive neuronal and glial cell death.^6^ Excessive glutamate release produces excitotoxicity and sustained calcium influx into neurons and oligodendrocytes, generating free radicals that damage cellular membranes and organelles and that disrupt cellular homeostasis through necrosis and necroptosis. ^7^ Ischemic injury to spinal cord white matter impairs action potential conduction, contributing to neurogenic shock and functional deficits.^8,9^ In the subacute and chronic phases, oligodendrocyte apoptosis and degeneration extend damage beyond the initial site of trauma, producing progressive demyelination and neurogenic pain.^10,11^ Effective therapeutic intervention must therefore mitigate both primary and secondary injury while supporting neuronal survival, remyelination, and functional recovery.

A promising neuroprotective strategy targets oxidative stress and inflammation, two convergent drivers of secondary damage. Curcumin, a naturally occurring polyphenol from the turmeric plant, has long attracted attention for its potent antioxidant, anti-inflammatory, and neuroprotective properties.^12,13^ It modulates numerous cellular signaling pathways implicated in disease (*e.g.* NF-κB, TNF-α, IL-6, IL-8, mTOR, MAPKs, and STAT) while upregulating NRF2, a master regulator of antioxidant defenses.^12^ Its neuroprotective potential is further supported by its ability to reduce nitric oxide production, inhibit lipid peroxidation, and suppress pro-inflammatory cytokines such as IL-1β and TNF-α in macrophages,^14–16^ to attenuate cerebral hypoperfusion-induced injury in animal models,^17–20^ and to protect against excitotoxicity through modulation of neurotrophic factors and NMDA receptor subunits.^21,22^ Despite these promising pharmacological effects, unmodified curcumin suffers from poor bioavailability, low aqueous solubility, rapid metabolism, and poor systemic absorption.^23–25^ The conventional response to this problem has been to adjust the formulation with various excipients for systemic administration: nanoparticles, liposomes, micelles, and physically loaded hydrogels have all been used to enhance curcumin solubility and stability.^24^ These approaches treat the polymer as a carrier and curcumin as a noncovalent payload. They suffer from rapid clearance by the mononuclear phagocyte system, unpredictable biodistribution, potential cytotoxicity at high concentrations, and release kinetics governed by carrier degradation or diffusion rather than by the timescale of secondary injury. For SCI specifically, none of these modalities solves the fundamental problem of sustaining therapeutic curcumin concentration, locally at the lesion site, over the course of the multi-week injury evolution window.

To fill that gap, we recently developed a fundamentally different approach in which drug molecules are not encapsulated in a carrier material, but rather covalently polymerized into the backbone of a macromolecular framework, yielding locally implantable biomaterials that serve as a depot of enchained curcumin pro-drug.^26–30^ In the present study, curcumin is copolymerized with poly(ethylene glycol) (PEG) and sebacoyl chloride to produce a polyester whose backbone alternates the antioxidant unit with PEG segments (Figure 1). Whereas free curcumin is unstable in aqueous media on the timescale of a few minutes, poly(curcumin-co-PEG) biomaterials are long-lived: they quench ROS on contact even after incubation in physiological buffer for more than two months. Copolymerization of the rigid/hydrophobic curcumin with the flexible/hydrophilic PEG produces a hard/soft dichotomy that enables tunable thermomechanical properties, hydrophilicity/solubility, and degradation rate as a function of curcumin:PEG ratio. The 50:50 composition (hence denoted as “P50”) used here represents a careful compromise: viscoelastic mechanical properties (G′ = 0.2 MPa ≫ G″ = 0.04 MPa at 37 °C), moderate hydrophilicity (θ ∼ 47°), and high molecular weight (∼18.6 kDa, Ð ∼ 3.3) that enables solution casting into free-standing thin films. P50 rescues neurons in culture from oxidative insult (100 µM H_2_O_2_) and shows excellent *in vitro* biocompatibility. As a structurally matched negative control we prepared the same polyester architecture in which curcumin is replaced with bisphenol A, a monomer of similar reactivity but no ability to quench reactive oxygen species. Because SCI patients in the emergency-care setting routinely undergo surgery to decompress spinal tissues and remove bone fragments,^31^ there is a clear clinical opportunity to implant a neuroprotective thin film on the dura mater during that same procedure, for maximal benefit and minimal added procedural burden.

**Figure 1.**
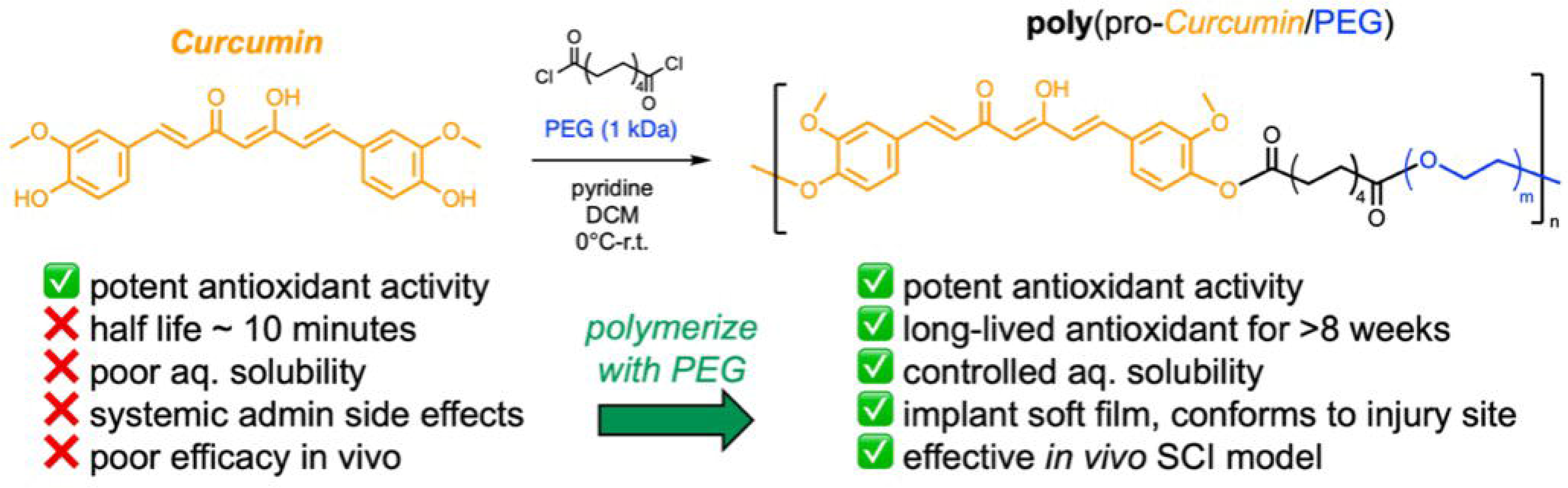
Synthesis of a 50:50 (mol %) copolymer containing curcumin and PEG with a sebacoyl chloride linker, dubbed “**P_50_**”. The obtained polyesters have tunable physical and thermomechanical properties, as well as vastly extended timescale of antioxidant activity.

In this study, we report the first example of a poly(pro-drug) thin film implanted locally at the injury site to enhance recovery for extended duration following SCI. Unlike conventional curcumin delivery strategies (liposomes, micelles, nanoparticles, or physically loaded hydrogels) in which curcumin is a noncovalent payload that diffuses or is released by carrier degradation, our architecture incorporates curcumin as a backbone repeat unit of the polymer itself. Release therefore requires slow cleavage of polymer ester bonds, which proceeds on a timescale matched to the multi-week duration of secondary injury rather than the minutes-to-hours timescale that governs aqueous curcumin or its conventionally analogs. The poly(curcumin-co-PEG sebacate) material used in this work has a molar ratio of curcumin:PEG of ∼50:50 and is thus denoted as “P50”. We prepare thin films by solution casting into ∼100 µm free-standing implants that are sufficiently rigid at room temperature (22-24 °C) for surgical handling yet soften to a tissue-compatible modulus, in the kPa range, upon implantation at 37 °C. Such tailored physical properties are tunable molecular design parameters rather than incidental outcomes of synthesis. Because the architecture is modular, the same design principle can be broadly utilized with any other phenolic and/or polyol-containing bioactives, opening a route to sustained local CNS delivery of neuroprotective, anti-inflammatory, or trophic small molecules beyond curcumin. We hypothesized that a single epidural implantation of a P50 film immediately after SCI^31^ would attenuate the secondary injury cascade and improve locomotor recovery, and tested this hypothesis in a moderate-to-severe T9 contusion model in mice, using myelin staining, immunohistochemistry of glial activation markers, and bulk RNA sequencing of tissue rostral and caudal to the lesion to dissect where and how the film acts.

## METHODS

### Poly(pro-curcumin) Thin Films

Synthesis of the statistical copolymer containing 50 mol% of curcumin and 50 mol% PEG in the feed (denoted as compound **P_50_**) was performed according to our previously published procedure,^26^ and gave similar results. The batch utilized in this paper is a copolymer with a peak MW of 18.6 kDa and a disperity of Đ∼3.3, according to analysis by GPC (see supporting info). The ^1^H NMR spectrum is consistent with our previously published work and gives an experimental copolymer composition of ∼48 mol% curcumin, relative to PEG. Films of **P_50_** were solution cast as previously published with minor modifications: briefly, a solution of **P_50_** in chloroform (8 wt%, 130 mg/mL) was prepared in a 20 mL scintillation vial, vortexed, and heated gently with a handheld heat gun until an optically clear yellow/orange solution was obtained. An aliquot of this solution (∼0.1 mL) was then carefully spread on to a clean glass microscope slide, and the solvent was allowed to slowly evaporate overnight in a fume hood. After 18 hours, the films were dried under vacuum (35 mTorr) overnight and manually cut into 2x4 mm^2^ nominal rectangles using a razor blade, and then gently lifted off the surface with a pair of fine-tipped forceps.

### Animal Model and Ethical Considerations

All animal procedures were conducted in accordance with the Public Health Service (PHS) Policy on Humane Care and Use of Laboratory Animals and the NIH Guide for the Care and Use of Laboratory Animals. Experimental protocols were approved by the Institutional Animal Care and Use Committee (IACUC) at the James J. Peters Veterans Affairs Medical Center (JJP VAMC) under protocol #CAR-20-11. Three-month-old male and female C57BL/6 mice were purchased from Charles River Laboratories and acclimated for one week prior to surgery. Mice were housed in a temperature-controlled environment (23-25°C) with a 12:12 h light/dark cycle (lights off from 6:00 AM to 6:00 PM) and provided ad libitum access to standard pelleted chow and water. To minimize potential social stress and aggression, animals were singly housed one week before surgery.

### SCI Surgery and P_50_ Film Implantation

At 18 weeks of age, mice of comparable body weights were randomly assigned to one of four experimental groups: (1) SCI + **P_50_**film, (2) SCI + control film (vehicle), (3) sham-SCI + **P_50_** film, and (4) sham-SCI + control polymer film. SCI was generated using the Infinite Horizons (IH) impactor (Precision Systems and Instrumentation) to generate a motor-incomplete contusion injury, as previously described.^32–35^ Briefly, mice were anesthetized with 3% isoflurane and maintained on a heating pad at 37°C. After hair removal and aseptic preparation, a laminectomy was performed at thoracic vertebrae 9 (T9) to expose the dura, creating a laminectomy window approximately 2 mm in diameter. Sham-operated animals underwent laminectomy without impact and had the paraspinal musculature sutured for spinal column stabilization before wound closure. For SCI groups, animals were positioned on the IH impactor platform, and the vertebral column was stabilized using forceps attached to the clamping system. A 65-kdyne contusion impact was applied, generating a moderate-to-severe injury known to produce partial locomotor recovery with a plateau at approximately 28 days post-injury. Left-right symmetry of the contusion was verified by uniform bruising at the dorsal median sulcus. Actual impact force and spinal cord displacement were recorded for each animal (see Supporting information, Figures S2 and S3). Immediately post-injury, either a poly-curcumin film or poly-control film was placed directly over the exposed dura. The films were secured in place by suturing the overlying musculature with resorbable sutures, followed by skin closure with 7-mm wound clips.

### Post-Operative Care

Animals were placed in clean cages with Alpha-Dri bedding (Newco Distributors, Inc., Hayward, CA, USA). Cages were placed on heating pads maintained at 37°C for the first 72 hours. All animals received pre-warmed lactated Ringer’s solution (LRS), carprofen (5 mg/kg), and enrofloxacin (Baytril, 5 mg/kg) once daily for five days. Bladders were manually expressed twice daily until spontaneous voiding resumed. Mice were monitored daily for signs of stress, urine scalding, autophagia, and general health status. Wound clips were removed at 10 days post-injury.

### Behavioral Testing

Hindlimb locomotor function was assessed at 3-, 7-, and 14-days post-injury using the Basso Mouse Scale (BMS) open-field locomotion test.^36^ The BMS is a validated 10-point scale that quantifies locomotor recovery, with 0 indicating complete paralysis and 9 representing normal gait.^36,37^ Mice were placed in an open-field arena and allowed to move freely. Two independent observers, blinded to treatment groups, scored locomotion based on key gait parameters, including paw placement, weight support, trunk stability, and limb coordination.

### Tissue Processing and Immunohistochemistry

Spinal cords were harvested after perfusion-fixation at endpoint and post-fixed in 4% paraformaldehyde for 24 hours before cryoprotection in graded sucrose solutions (10%, 20%, and 30%). A 2-mm spinal cord segment, spanning 1 mm rostral and 1 mm caudal to the lesion epicenter, was dissected, bisected at the lesion core, and embedded in optimal cutting temperature (OCT) compound. Transverse sections (10 μm) were obtained using a cryostat (Leica Microsystems, West Hollywood, CA, USA) and mounted for immunohistochemistry. Myelin integrity was assessed using FluoroMyelin Green (Invitrogen, F34651). Sections were permeabilized in Triton X-100, incubated with FluoroMyelin for 20 minutes, washed, and imaged using a confocal microscope (Carl Zeiss, Jena, Germany). To visualize differences in spared myelin an image analysis was performed using the ImageJ software as described elsewhere (Streijger, Lee et al. 2016). Briefly, thresholds for signals in the images were set at background gray values. The region of interest was determined and mean intensity and total number of pixels above threshold were measured. Spared tissue was determining by the percentage of white matter calculated by the total area of the spinal cord for each section using ImageJ. Glial cell activation was evaluated by immunostaining for glial fibrillary acidic protein (GFAP; Abcam #ab7260 at 1:500 dilution) to identify reactive astrocytes and ionized calcium-binding adapter molecule 1 (Iba1; Abcam #ab178846 at 1:500 dilution) to detect activated microglia/macrophages. Sections cut 400 μm rostral and caudal to the lesion were permeabilized, stained, and imaged by confocal microscopy. Secondary antibodies Alexa Fluor 488-conjugated goat anti-mouse IgG (Abcam # ab150113) and Alexa Fluor 647-conjugated goat anti-rabbit IgG (Abcam # ab150079) were used both at 1:2500 dilution. 5 x 5 tiled images were obtained from stained sections using a 20X objective and a Zeiss 700 confocal microscope (Carl Zeiss). Blinded quantification was performed using ImageJ software (version 2.1.0/1.53c, National Institute of Health, USA) and integrated density of pixels was measured for each section and a mean value was calculated as previously described (Oliveira, Thams et al. 2004, Toro, Hansen et al. 2021). Maximum background threshold was determined for each image and set for intensity quantification. Data are represented using the mean ±SEM.

### RNA Sequencing

Total RNA was extracted from spinal cord tissue segments located immediately rostral and caudal to the injury epicenter using a TRIzol-based protocol as previously described (Toro, 2021; Toro, 2023; Toro, 2024). RNA integrity was assessed using the Agilent 4200 TapeStation system and samples with RNA integrity numbers (RIN) >7.0 were used for downstream processing. Library preparation was performed using the SMARTer Stranded Total RNA-Seq Kit v2 (Takara), and sequencing was conducted on the Illumina NextSeq 2000 platform using paired-end 100 cycle protocol. Raw sequencing data (FASTQ files) were pre-processed with HTStream to remove adapter sequences and low-quality reads. The first three nucleotides of the second sequencing read were trimmed using the seqtk tool according to the SMARTer kit instructions. The reads were then aligned using software STAR (v. 2.7.11b) to the reference mouse genome (GRCm39) with GTF transcript annotations (gencode.vM38), and gene-level counts were quantified using feature Counts (subread R package, v. 2.0.1) Differential gene expression analysis was performed using DESeq2 (v. 1.44.0). Gene ontology (GO) analysis was performed with an online GO tool WebGestalt (www.webgestalt.org).

## RESULTS and DISCUSSION

### Poly(pro-Curcumin/PEG) Film Casting

Synthesis of the **P_50_** raw material was carried out according to our prior report^26^ and characterization by ^1^H NMR and GPC confirm the proposed structure was indeed obtained. Viscoelastic thin films of polymerized curcumin material **P_50_** were then processed by solvent casting according to the procedure described above. We obtained rectangular films with experimental dimensions of 1.7 ±0.2 × 3.3 ±0.3 mm, averaged over a total of 56 samples cast (Figure 3A). The thickness is consistent and smooth at ∼100 µm by profilometry (Figure 3B). The sample mass is nominally ∼0.5 mg total polymer. Immediately following casting, the film has a consistency of a viscous semisolid and was not conducive to lifting a free-standing film that could be manipulated by a surgeon. However, physical aging of the films (stored at room temperature, protected from light, for at least 2 weeks and up to 4 months) successfully enables densification and improved mechanical properties, such that the films can be delaminated from the glass slide with a razor blade and lifted with fine-tipped forceps for implantation. The films also become noticeably more deep orange compared to the light-yellow color upon initial casting. Such a color change is indicative of curcumin π-π stacking in the solid state,^38^ which can behave as physical crosslinks. In order to explain the molecular basis of the physical ageing process, we performed differential scanning calorimetry (DSC) on freshly cast versus aged films (Figure 2D). The fresh sample shows a glass transition temperature (T_g_) around -43 °C, and no observable melting endotherm, which is in good agreement with our published report.^26^ We found that the position of the glass transition (T_g_) is largely unchanged upon aging, but the aged sample demonstrated the emergence of a significant crystalline melting endotherm (T_m_) around 27 °C, which is comparable to reported melting temperatures for PEG.^39,40^ This result suggests slow room-temperature crystallization of the PEG segments is the root cause of physical aging. The T_m_ is just slightly above room temperature, suggesting that the aging process in ambient conditions yields the ideal annealing temperature to thermally induce crystallization. Thus, we conclude that the effect of physically aging films post-casting is primarily ascribed to the slow crystallization of the PEG component in the semisolid state. It is also noteworthy that the materials are just on the edge of solid-like properties at room temperature, which is highly beneficial for our intended application. At room temperature, they can be manipulated as free-standing solid films with fine-tipped forceps. When implanted at 37 °C, they will melt and soften considerably, with additional plasticization due to water uptake, and thus will possess excellent mechanical compatibility with the soft tissues of the CNS. Finally, as we previously showed, P50 is partially water soluble, such that a small amount of PEGylated curcumin elutes locally from the mostly insoluble films immediately after implantation, thus enabling a combination of immediate burst of antioxidant activity, combined with long-lived on-contact activity. As a control, we also repeated the characterization of the polymer films by ^1^H NMR and GPC, as described above for the raw polymer material as-synthesized, on both the as-cast and aged films. No changes are evident in the chemical structure by NMR (supporting information, Figure S1) nor the molecular weight distribution by GPC (Figure 3C), which confirms that the physical aging process does not involve any chemical alterations to the structure.

**Figure 2.**
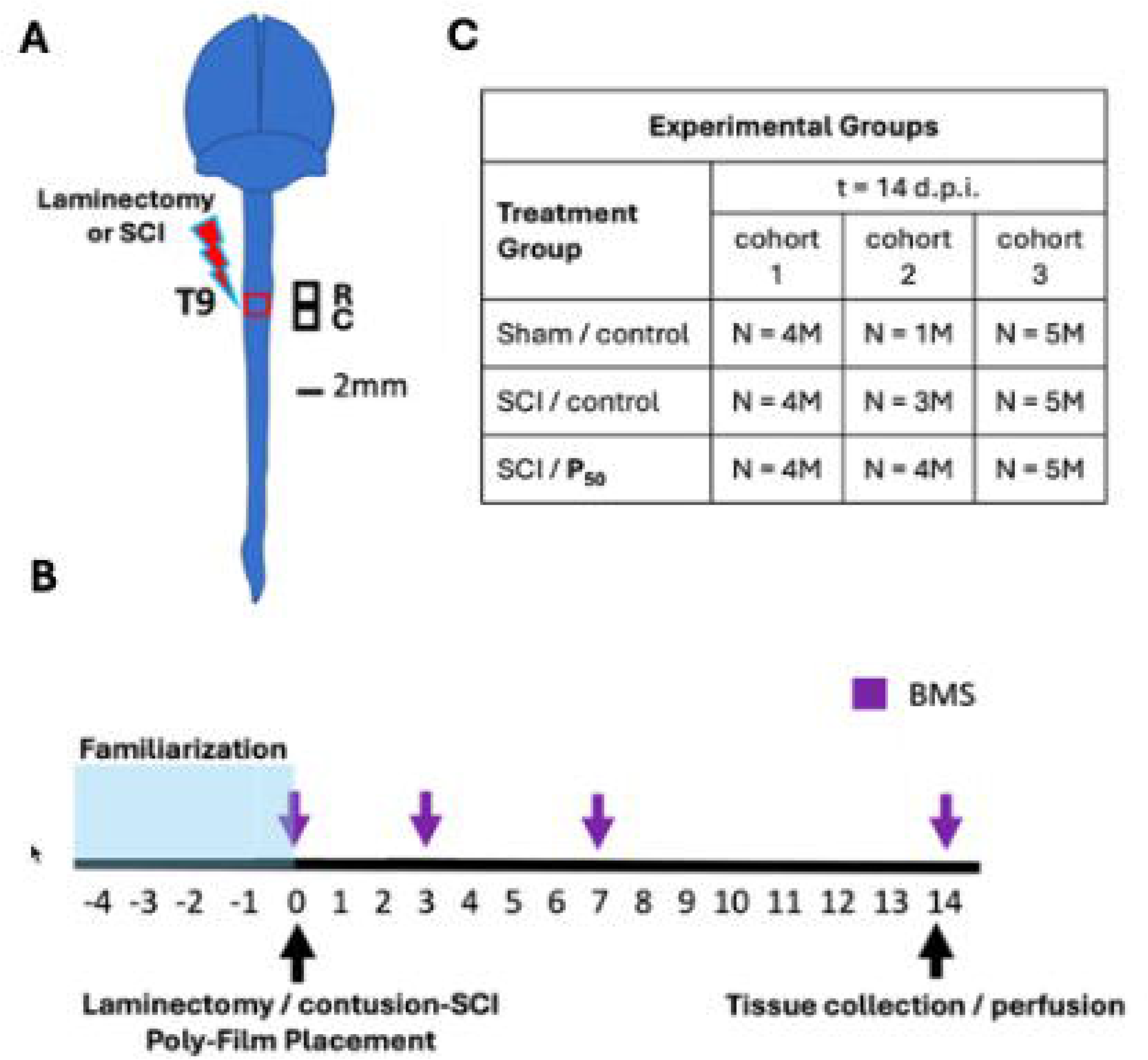
Experimental design. (A) Anatomical representation of the mouse brain and spinal cord showing the site of laminectomy or contusion SCI (red box) at the level of thoracic vertebrae 9 (T9). Black squares represent locations of the rostral and caudal segments of spinal cord collected for histology and RNA-seq studies. R□=□rostral; C□=□caudal. Scale bar, 2□mm. (B) Timeline for animal familiarization with handling, behavioral testing, and tissue collection/animal perfusion are depicted. BMS, Basso mouse scale. (C) Table showing experimental groups and animal numbers. M: males; dpi: days post injury

**Figure 3.**
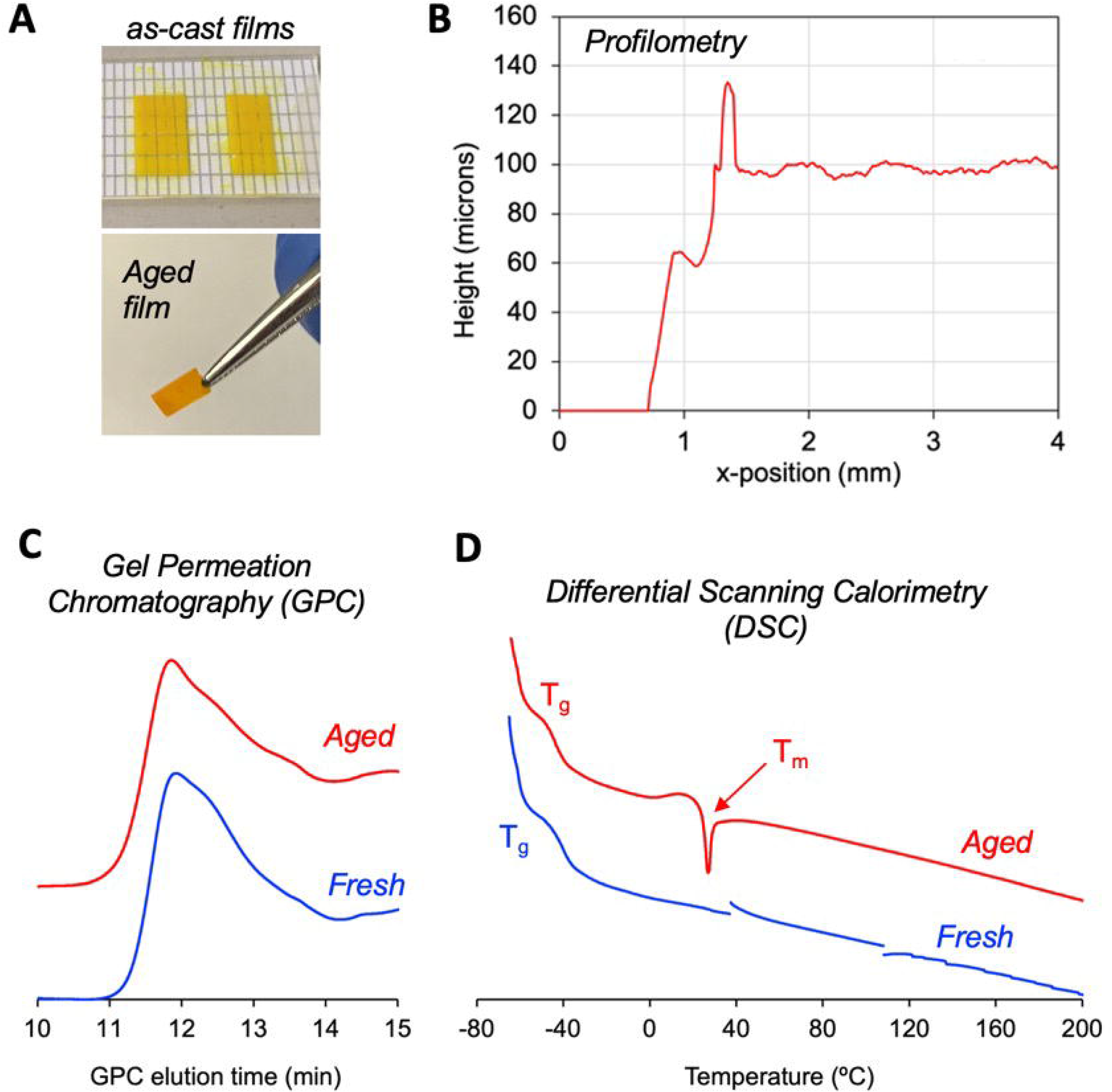
(A) Solvent cast thin films are prepared on glass slides. Following one month of storage under ambient temperature (protected from light), the films become robust enough to handle as free-standing thin films. (B) Profilometry reveals the films are smooth and have an average thickness of about 100 µm. (C) gel permeation chromatography (GPC) shows that the molecular weight distribution of **P_50_**does not substantially change upon aging (D) differential scanning calorimetry (DSC) shows that the PEG component of the copolymer crystallizes upon aging, which explains the improvement in mechanical robustness.

### Behavioral Testing

To evaluate the functional recovery of hindlimb locomotion following SCI, BMS scores were recorded at baseline (pre-surgery) and at 3-, 7-, and 14-days post-injury. Performance was compared to that of animals that underwent laminectomy without SCI, as “sham” (**Figure 4**). As expected, all animals that received SCI, regardless of treatment condition, exhibited a significant decline in BMS scores immediately following injury, with an average score of 0.44 ± 0.076 in the vehicle-treated group and 0.67 ± 0.19 in the **P_50_** treated group. There was no significant difference between the groups at day 3 post-injury (p = 0.6085), indicating that the severity of the initial injury was comparable across conditions. By day 7, locomotor recovery was observed in both SCI groups. However, animals treated with poly-pro-curcumin exhibited significantly greater functional improvement compared to vehicle-treated counterparts (p < 0.05). While vehicle-treated animals only displayed ankle movement, poly-pro-curcumin-treated animals were able to maintain weight support throughout an entire step. Functional improvement continued through day 14 post-injury, where poly-pro-curcumin-treated mice demonstrated a significantly higher mean BMS score (3.48 ± 0.19) than vehicle-treated animals (2.31 ± 0.21) (p < 0.01). These findings suggest that poly-pro-curcumin enhances early motor recovery following SCI.

**Figure 4.**
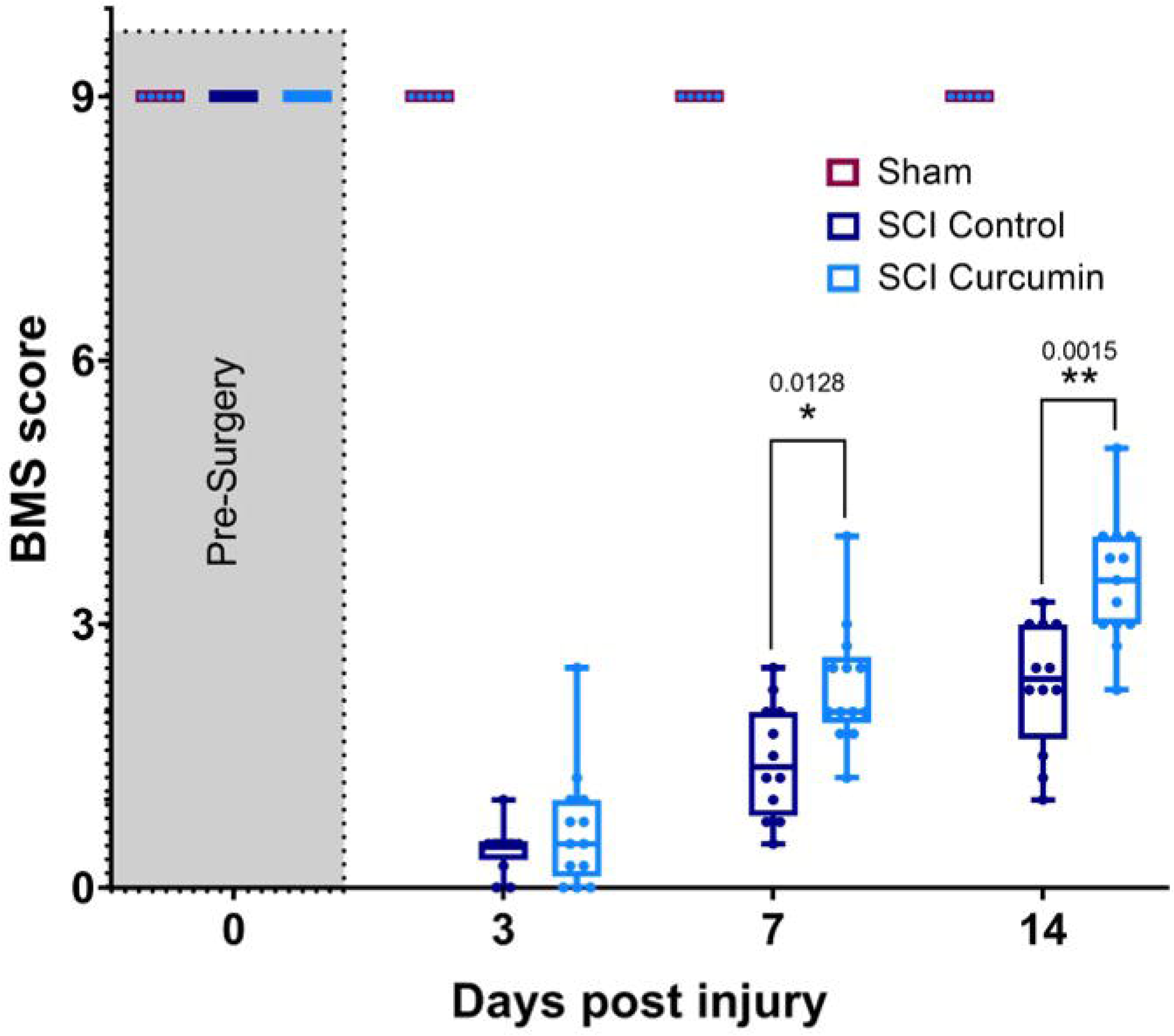
Basso Mouse Scale (BMS) scores were determined prior to surgery then repeated at the indicated times. Box-and-whisker diagrams represent the median, third quartile (upper edge) and first quartile (lower edge), and minimum and maximum values (whiskers) of the data. Data were analyzed by mixed model ANOVA using a Bonferroni’s test post-hoc. *, p < 0.05; **, p < 0.01; ***, p < 0.001.

Functional locomotor improvements in poly(pro-curcumin)-treated animals, while modest, were statistically significant, indicating that early intervention with this compound may facilitate meaningful motor recovery. An increase in BMS score from 2 to 3 corresponds to a transition from ‘extensive ankle movement’ to ‘plantar placing of the paw’ and even ‘occasional dorsal stepping with weight support’ - a substantial improvement in locomotion. Translating this into a clinical context, such a change could represent the difference between minimal voluntary leg movement and the ability to ambulate with assistive devices, significantly improving quality of life. The benefits for individuals with incomplete SCI may exceed those seen here, underscoring the importance of evaluating poly-pro-curcumin across different injury severities. These results align with previous findings that suggest curcumin and its derivatives may enhance neuroprotection and functional recovery in CNS injuries.

### FluoroMyelin Staining and Myelin Preservation

To evaluate myelin integrity following injury, spinal cord sections were stained with FluoroMyelin and analyzed at 200 µm intervals rostral and caudal to the lesion site (**Figure 5**). Quantification of myelinated area, analyzed via two-way ANOVA, revealed a significant main effect of poly-pro-curcumin treatment (p < 0.05), accounting for approximately 17% of the total variance. Moreover, post-hoc comparisons reveal significant differences in myelination at 50 µm distance rostral and caudal from the lesion epicenter. However, no significant differences were detected elsewhere between rostral and caudal regions. These findings suggest that the curcumin-containing **P_50_** film implant preserves myelin or promotes remyelination after SCI, potentially by mitigating oligodendrocyte loss or enhancing repair mechanisms. The observed increase in myelination is consistent with the improved BMS scores in poly-pro-curcumin-treated animals, supporting the hypothesis that polycurcumin films increased the number of spared myelinated fibers. Notably, white matter preservation was significantly greater in polymerized curcumin **P_50_**-treated animals compared to controls. This finding suggests that curcumin may exert a neuroprotective effect by mitigating oligodendrocyte loss, reducing excitotoxic damage, or promoting remyelination, mechanisms that warrant further exploration. White matter integrity is crucial for signal transmission across the spinal cord, and its preservation strongly correlates with better functional outcomes after SCI.^41–43^

**Figure 5.**
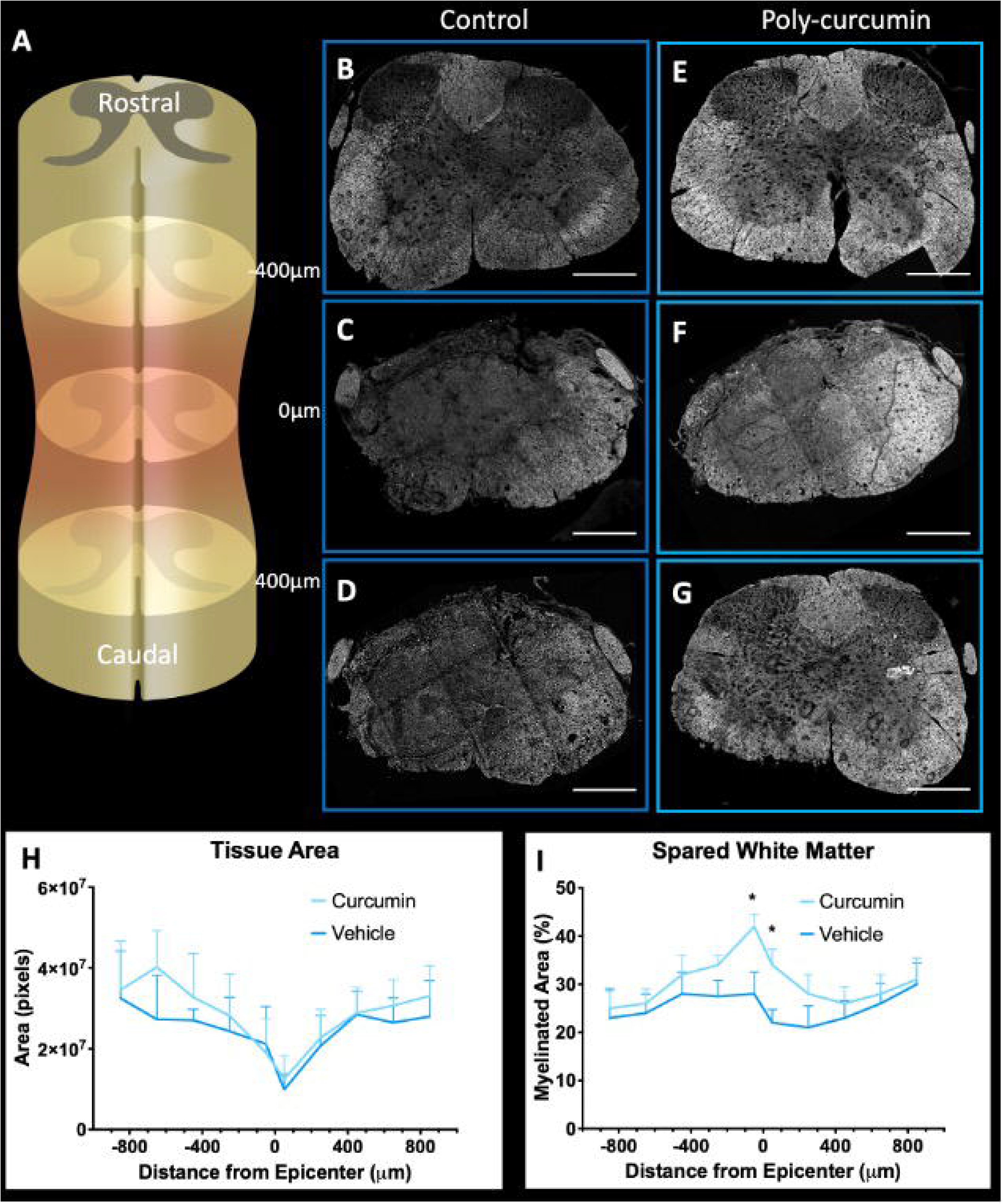
Poly(curcumin) films promote sparing of white matter in mice after SCI. (A) Scheme of a mouse spinal cord showing areas rostral, caudal of the injury epicenter. (B-G) Perfusion-fixed spinal cords of control (B-D) or poly(curcumin) (E-G) implanted films following SCI were cryo-sectioned at 14 days. Transverse sections were collected and stained with FluoroMyelin. Representative images at 400 μm rostral from the injury epicenter (B, E), at the epicenter (C, F), and 400 μm caudal from the epicenter (D, G) are show . Tissue area (H) and fluorescence intensity (I) was quantified for each of 4 different animals per group. \**p* < 0.05; unpaired Student’s *t*-test. Scale bar:

### Immunohistochemical Analysis

To assess cellular and tissue-level changes following SCI, spinal cord sections spanning the lesion site were immunohistochemically stained for markers of gliosis. Astrocyte reactivity was examined via GFAP staining, which revealed prominent astrocytic activation surrounding the lesion in both poly-pro-curcumin- and vehicle-treated animals. As expected, astrocytes formed a dense, scar-like boundary around the lesion core (**Figure 6**), consistent with previous reports of glial scar formation post-SCI.^44^ GFAP expression appeared lower in each sample of the poly-pro-curcumin group. Moreover, quantitative analysis revealed a statistically significant difference between groups rostral from the injury site (p<0.0486).

**Figure 6.**
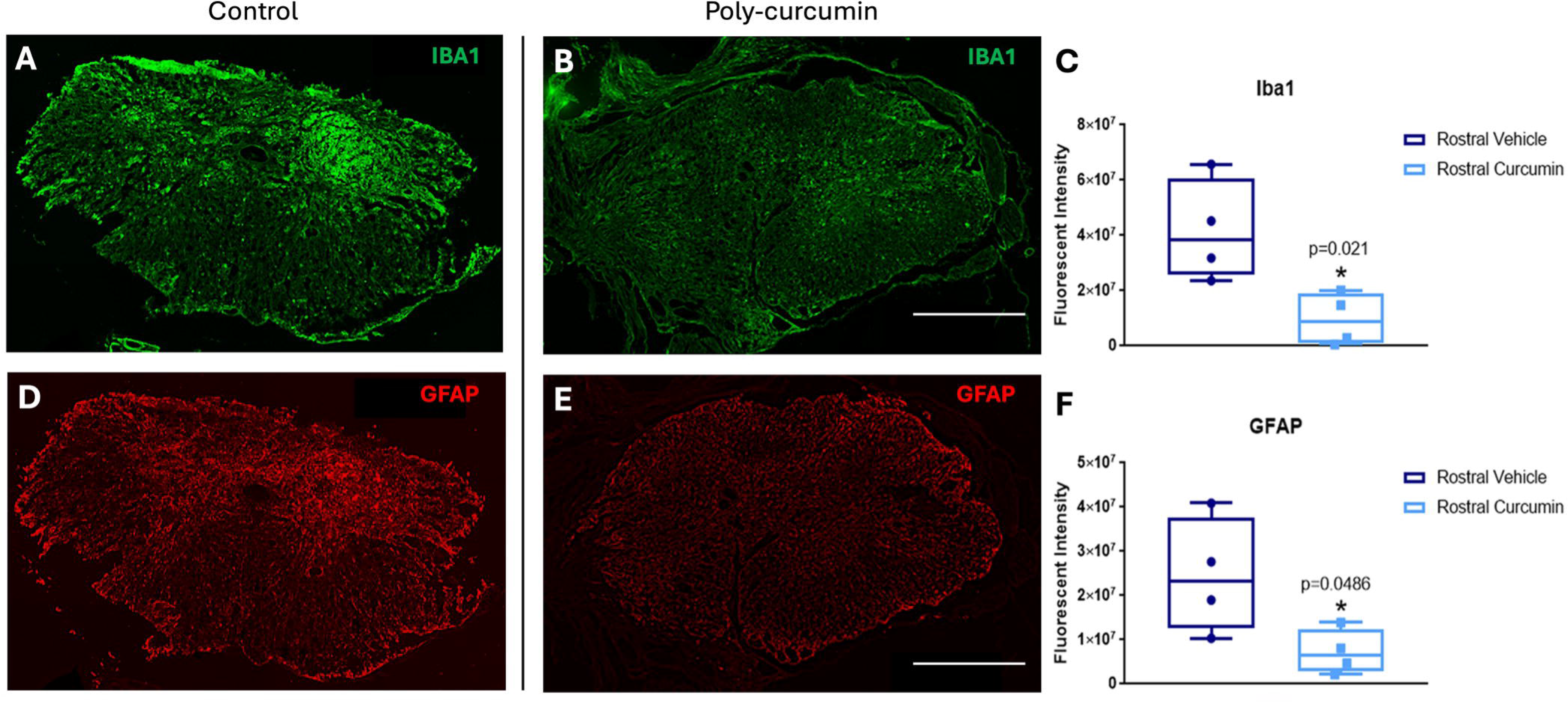
Poly(curcumin) films decrease glial response in mice at 14 days after SCI. Transverse spinal cord sections taken 200 µm rostral and caudal to the SCI lesion were immunostained and imaged using confocal microscopy. Sections from SCI animals with control films and poly-curcumin films were labeled for Iba1 (A, B; green) and GFAP (D, E; red). Quantification of staining intensity of Iba1 (C) and GFAP (F) was performed for four animals per group. Box-and-whisker diagrams represent the median, third quartile (upper edge) and first quartile (lower edge), and minimum and maximum values (whiskers) of the data. *p < 0.05; unpaired Student’s t-test comparisons. Scale bar: 500 μm.

Microglial/macrophage activation was assessed using Iba1 staining. Iba1-positive cells were distributed throughout the injured spinal cord, with a higher density in vehicle-treated animals. Similar to astrocytic reactivity, quantitative analysis indicated a significant reduction of Iba1 expression in poly-pro-curcumin-treated mice, (p <0.021). These findings support the idea that poly-pro-curcumin attenuate glial response following SCI. Additional studies are needed to confirm these effects with higher statistical power.

Glial scar plays a dual role in SCI recovery: it protects some tissue from further damage but also creates a physical and biochemical barrier that impedes neural regeneration. A reduction in glial scar formation in **P_50_**-treated animals may suggest an environment more conducive to repair and functional recovery.

### Bulk Transcriptomics

To investigate transcriptomic changes associated with poly-pro-curcumin treatment, bulk RNA sequencing (RNA-seq) was performed on spinal cord tissue samples obtained from regions immediately rostral and caudal to the lesion epicenter (N=4-5 samples for each treatment condition and tissue region). In rostral samples, we detected 39 differentially expressed genes (DEGs) using a fold-change threshold of >20% and an FDR-adjusted p-value < 0.10 (Benjamini– Hochberg), and 170 DEGs with a threshold of adjusted p-value < 0.20 (**Suppl. Table 1**). The distribution of curcumin-upregulated and downregulated genes (**Figure 7**) showed that the majority of DEGs were downregulated in polycurcumin film SCI animals. Notably, in contrast to rostral samples, no DEGs were detected in caudal samples at any significance threshold (**Suppl. Table 2**). We performed a functional analysis of DEGs (adjusted p-value <0.20) from rostral tissue using a gene ontology (GO) analysis tool WebGestalt, and detected multiple significantly enriched GO terms, including those associated with inflammation or immune response (e.g., “*neuroinflmammatory response*”, “*chemokine production*”, “*interleukin-6 production*”, “*leukocyte proliferation*”) (**Figure 8**, and **Suppl. File 1**). Similar results (e.g., “*interleukin-17 production*” and “*leukocyte proliferation*”) with a smaller number of significant GO terms were obtained for DEGs with adjusted p-value <0.1 (data not shown).

**Figure 7.**
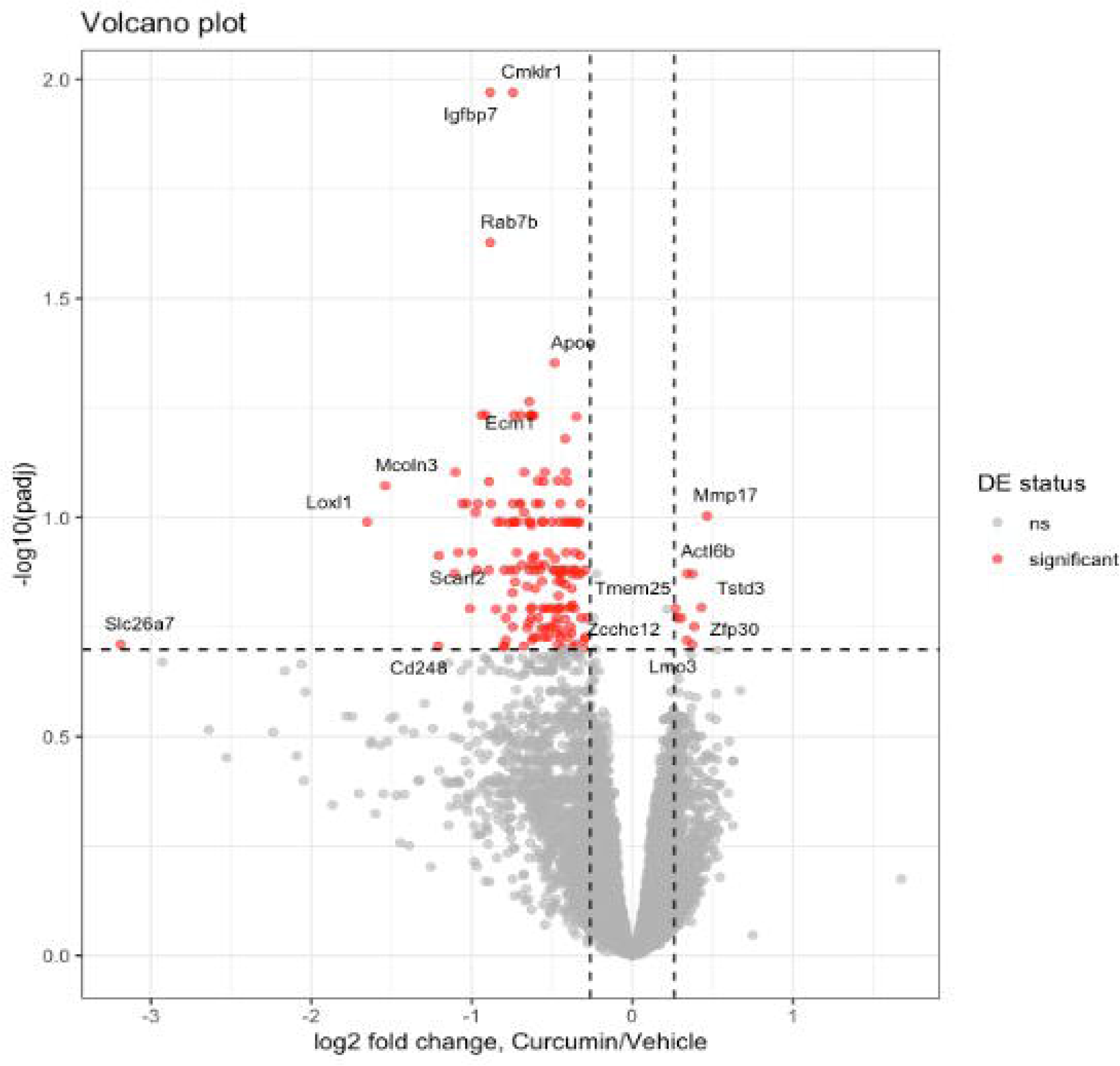
Volcano plot illustrating the direction and magnitude of expression change in DEGs from tissue samples rostral to the CI lesion. padj: FDR-adjusted p-value. DE status of genes: red – significant, gray - non-significant (“ns”). Thresholds to define significant DEGs: FDR-adjusted p-value < 0.2, fold change > 20%.

**Figure 8.**
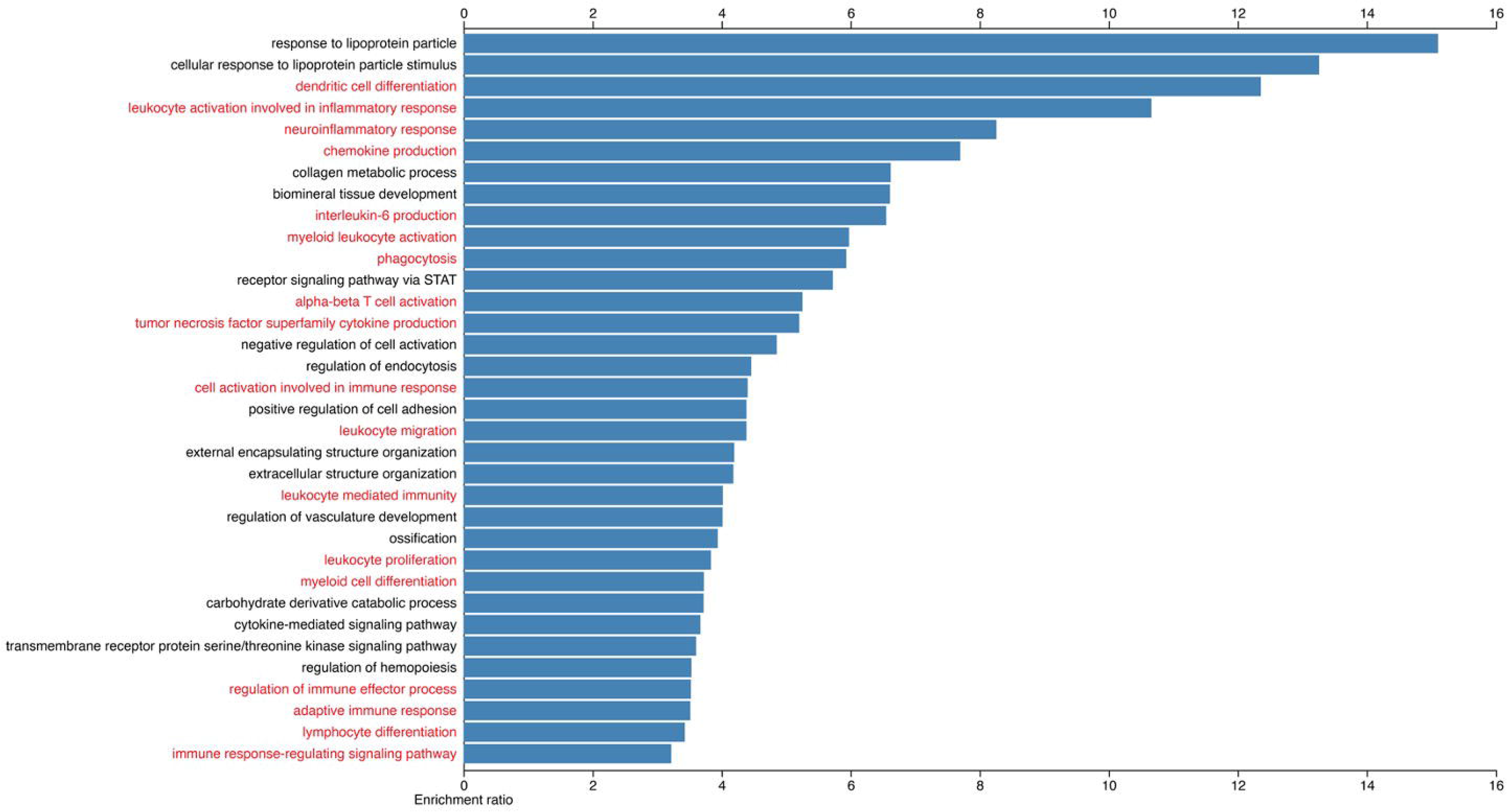
Gene Ontology (GO) analysis of rostral DEGs (FDR < 0.2, fold change > 20%) using a GO analysis tool WebGestalt. Shown are significantly enriched GO terms related to Biological Processes. GO terms associated with immune system function are highlighted in red.

Thus, at the molecular level, bulk RNA sequencing data showed downregulation of genes and pathways associated with inflammation in **P_50_**-treated animals compared to vehicle-treated controls at 14 days post injury. This result confirms our expectations based on the putative ability of curcumin, and polymers thereof, to quench reactive oxygen species.^45^ In fact, studies which compare curcumin to corticosteroids, in terms of SCI recovery potential, have found more benefit from curcumin.^46^ For example, curcumin in DMSO administered to rats reduced lipid peroxidation as compared to methylprednisolone sodium succinate.^47^ Future analyses will focus on identifying specific regulatory networks that mediate curcumin’s well-known neuroprotective effects, providing insight into potential combinatorial treatment strategies. Further, these results demonstrate the extended timescale of protection afforded by the polymeric viscoelastic form of curcumin, as opposed to the short-lived small molecule itself, considering that these changes in gene expression are observed 14 days post-injury.

## CONCLUSIONS

In this study, we investigated the effects of an epidural thin film biomaterial implant comprised of a copolyester derived from curcumin and PEG sebacate (**P_50_**), applied immediately after traumatic contusive SCI in mice. Through behavioral, histological, immunohistochemical, and transcriptomic analyses, we show that poly(curcumin)-based films attenuate key components of secondary injury, leading to improved functional and tissue-level outcomes. Specifically, **P_50_** films enhanced sensorimotor recovery, increased white matter sparing, and reduced activation of pro-inflammatory gene expression programs relative to control films. Together, these findings demonstrate that polymerized small-molecule drugs, or poly(pro-drugs), can be engineered into thin films that are both mechanically suitable for epidural implantation and biologically effective in modifying the post-injury milieu.

At the tissue level, **P_50_** treatment preserved myelinated white matter around the lesion epicenter and reduced indices of gliosis, as evidenced by decreased GFAP and Iba1 immunoreactivity rostral to the injury site. These changes likely contribute to the observed improvements in BMS scores, given the well-established relationship between white matter integrity, axonal conduction, and locomotor function after SCI. The rostro-caudal asymmetry in molecular and cellular effects marked downregulation of inflammatory pathways rostral, but not caudal, to the lesion suggests that poly(curcumin) films may preferentially modulate injury-propagating fronts and oxidative/inflammatory signaling gradients that extend from the primary site of impact. Bulk RNA sequencing further supports this interpretation by revealing coordinated downregulation of genes associated with neuroinflammatory and immune responses in **P_50_**-treated animals, consistent with sustained local quenching of reactive oxygen species and dampening of secondary inflammatory cascades.

These results also reinforce the broader concept that polymerizing bioactive small molecules into viscoelastic biomaterials can overcome key pharmacokinetic barriers that limit conventional systemic delivery. Curcumin’s clinical translation has historically been hindered by its poor solubility, rapid metabolism, and short systemic half-life; in contrast, embedding curcumin within a degradable polymer matrix provides a long-lived local reservoir that maintains antioxidant capacity over extended timescales while remaining confined to the injury site. In the context of SCI, where “time is spine” and early intervention can have durable effects on long-term outcome, an epidural thin-film approach is particularly attractive because it can, in principle, be integrated into standard decompression and stabilization surgeries without requiring repeated dosing or complex delivery hardware.

From a translational perspective, our data support the notion that epidural poly(pro-drug) films could serve as a modular platform for chronic, localized delivery of neuroprotective agents following SCI. The **P_50_** formulation evaluated here represents one instantiation of this platform, but the same design principles could be extended to other antioxidant, anti-inflammatory, or trophic small molecules, alone or in rational combination. Tunable polymer composition, degradation kinetics, and film geometry may allow tailoring of drug release profiles to different injury severities, anatomical levels, or clinical scenarios, including both acute traumatic injuries and chronic compressive myelopathies. Moreover, because these materials are applied epidurally, they avoid direct penetration of neural parenchyma, which may simplify regulatory and safety considerations.

Several limitations of the present work should be acknowledged. We examined a single injury severity and a single **P_50_** formulation and dose at an early post-injury time point, which constrains our ability to define dose–response relationships or long-term durability of benefit. The sample size, while sufficient to detect statistically significant changes in locomotor recovery and selected histological and transcriptomic endpoints, limits our power to detect more subtle effects, including sex-specific responses. We did not directly quantify in vivo degradation of the films, curcumin release kinetics, or tissue drug concentrations in the spinal cord, and therefore cannot yet link pharmacodynamics to polymer erosion profiles. In addition, our bulk RNA-seq data, while informative, do not resolve cell-type–specific transcriptional changes, which will be important for understanding how poly(curcumin) modulates distinct glial and neuronal populations.

Future studies should address these limitations by incorporating longitudinal behavioral and imaging assessments, multiple film compositions and doses, and extended follow-up to determine whether early gains in function are maintained or further amplified over time. Single-cell or spatial transcriptomic approaches will be valuable for mapping how poly(curcumin) reshapes cellular networks within and around the lesion. It will also be important to evaluate the compatibility of epidural poly(pro-drug) films with other emerging therapies, including rehabilitative training, cell or graft-based strategies, and systemic neuroprotective agents, to test for additive or synergistic effects.

In summary, our findings provide proof-of-concept that an epidural poly(curcumin-co-PEG) thin film can mitigate secondary injury mechanisms, preserve white matter, and improve early locomotor recovery after SCI in mice. These data position polymerized pro-drug thin films as a promising, adaptable platform for local pharmacologic intervention in SCI and potentially other CNS injuries, warranting continued preclinical development and careful consideration of their path toward clinical translation.

## FINANCIAL SUPPORT

NYS SCIRB Fellowship C3906GG to N.P.J; NYS SCIRB program support C38329GG; CSTA UL1TR004419 to M.M.S. & E.F.P.; CEPM 21-1216 to M.M.S.; United States Department of Veteran’s Affairs (I01RX003502-01A1), National Science Foundation (#2217513) E.F.P, and NYS SCRIB (C38335GG) to E.F.P. and R.J.G.; VA seqCURE program at JJP VAMC.

## AKNOWLEDGEMENTS

The authors would like to acknowledge the generous support of the Albany Stratton Veterans Affairs Medical Center, Albany, NY; James J Peters Veterans Affairs Medical Center, Bronx, NY; VA Sequencing Collaborations United for Research and Epidemiology Program (VA seqCURE), and the Bronx Veterans Medical Research Foundation, Bronx, NY.

---

**Figure S1.**
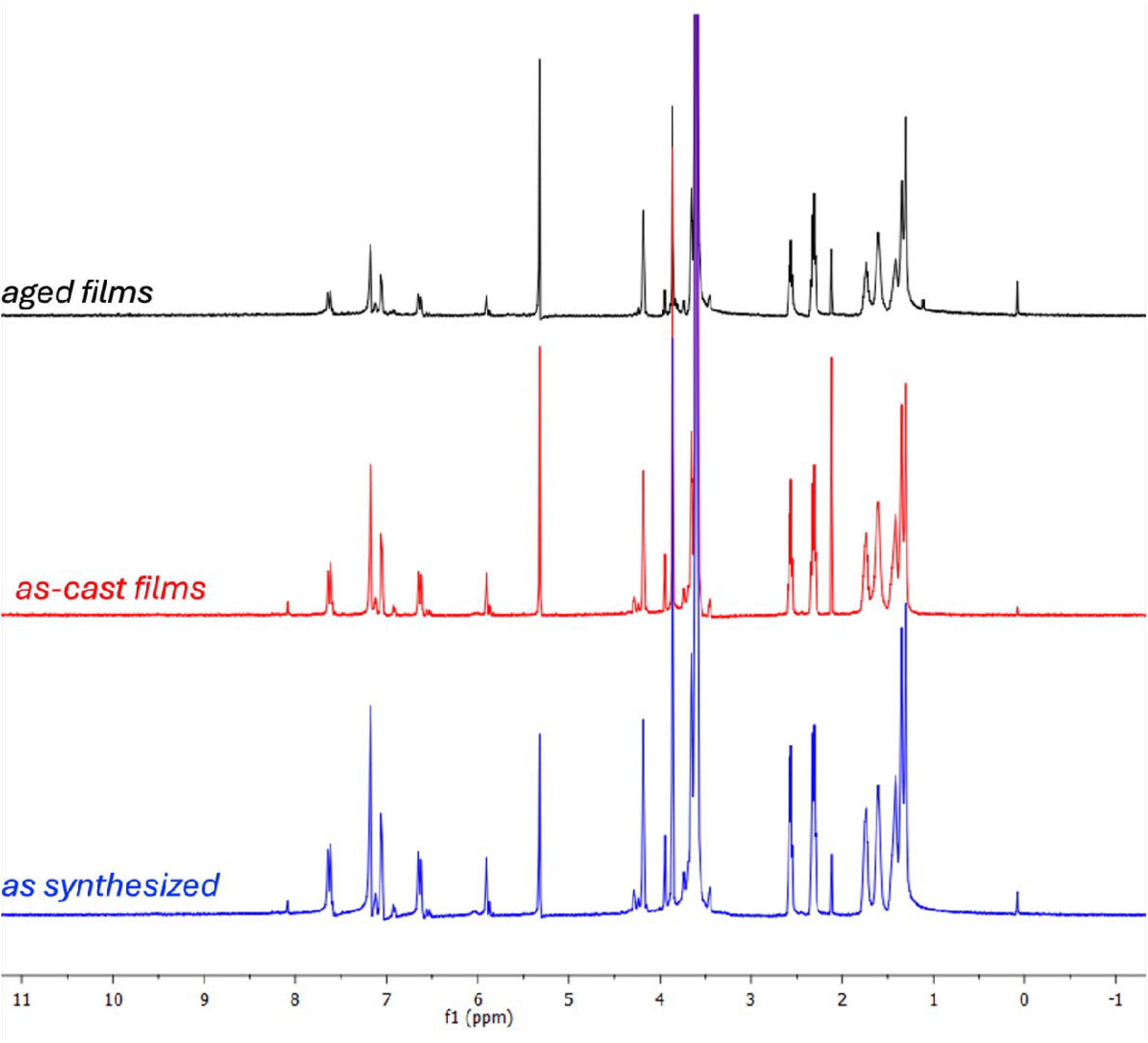
^1^H NMR spectra (CDCl_3_) of the P_50_ material as the raw powder from synthesis (blue), after film casting process (red), and after film casting and aging in air for 1 month (black). No change in chemical structure is evident, confirming that although physical aging improves mechanical properties, there is no degradation of chemistry when stored under ambient conditions (standard temperature, pressure, and relative humidity), as a dry solid film in the dark.

**Figure S2.**
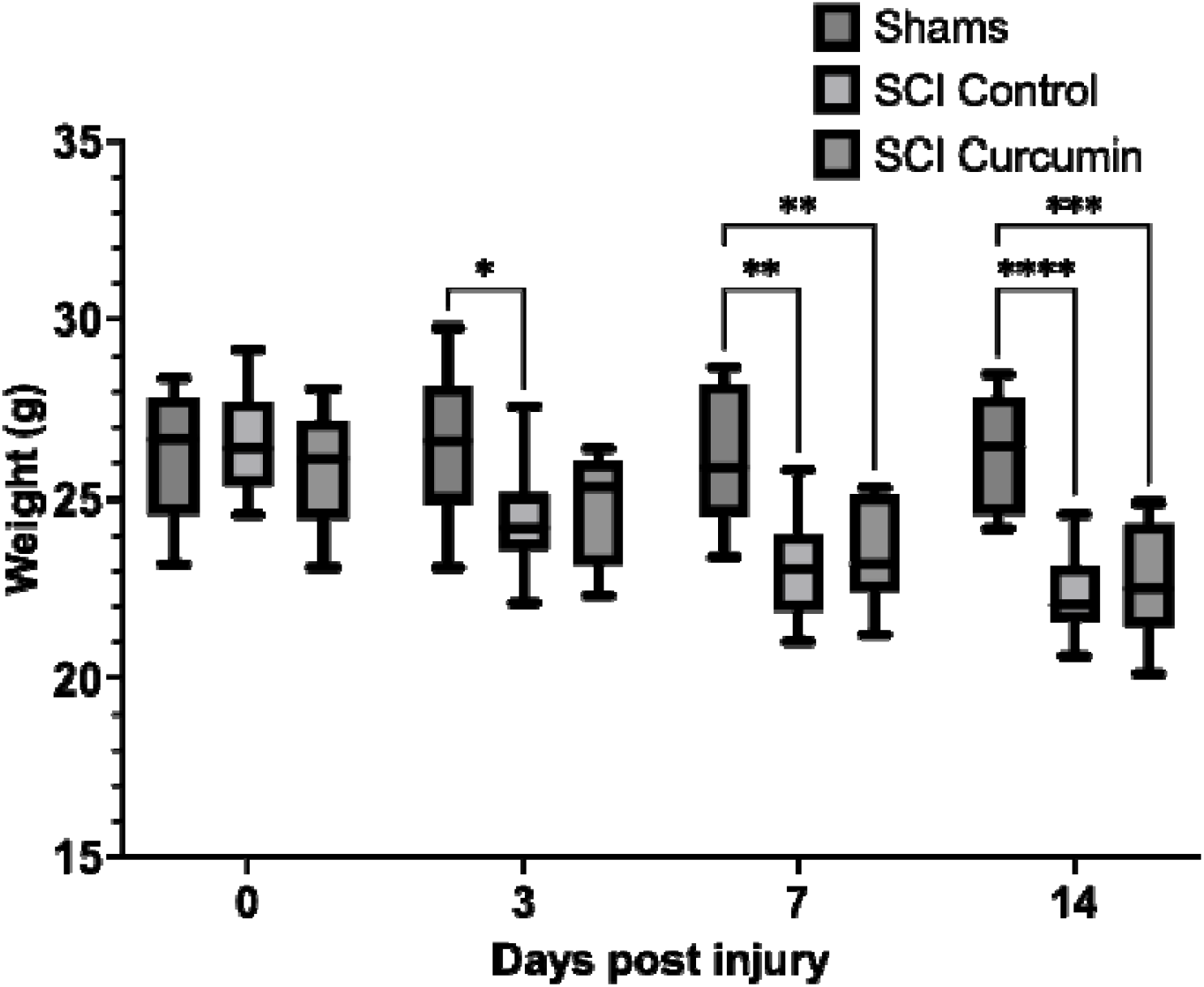
Weight loss as a function of time post-injury for shams, SCI with control film, and SCI with **P_50_**film implant.

**Figure S3.**
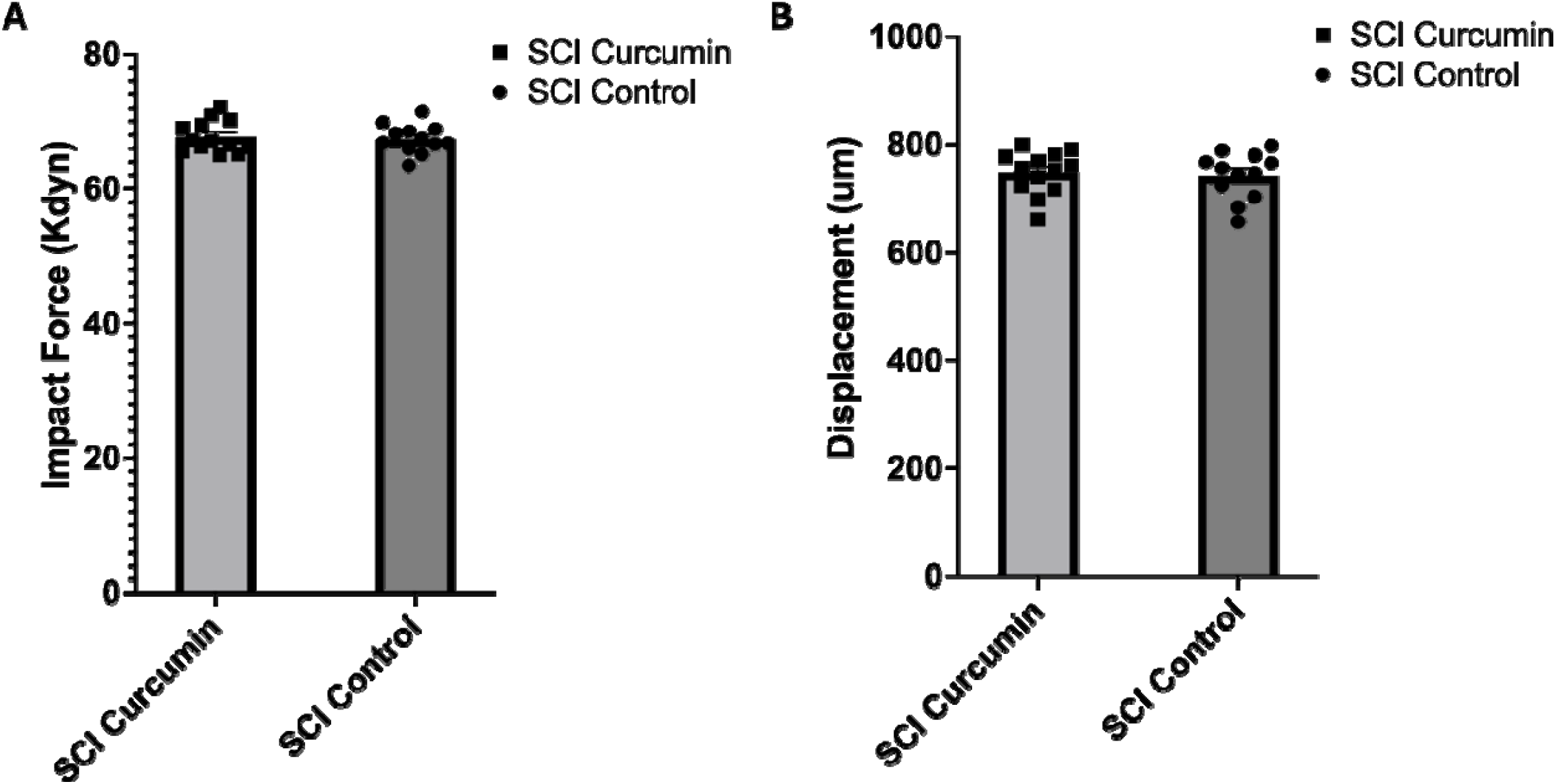
Measurement of impact force (A) and displacement (B) for the injuries sustained by mice with control film versus **P_50_** film implants show no difference in severity.

## Notes

### Competing Interest Statement

The authors have declared no competing interest.

### Summary of Updates

The manuscript has undergone extensive revision to improve clarity, scientific rigor, and overall impact. We strengthened the central rationale by more clearly defining the limitations of systemic pharmacotherapy for SCI and the need for sustained, lesion-localized therapeutic delivery. The Introduction and Discussion were revised to better position the poly(pro-drug) thin-film approach within the context of SCI neuroprotection and biomaterial-based drug delivery. Descriptions of film composition, epidural implantation, duration of activity, and the rationale for immediate post-injury placement were also refined. The in vivo experimental sections were strengthened through clearer descriptions of the T9 contusion SCI model, treatment and control groups, outcome measures, and statistical analyses. The presentation and interpretation of functional, histological, and molecular findings were revised to more clearly demonstrate our findings. Overall, these revisions substantially strengthen the manuscript and provide a clearer and more mechanistically grounded presentation of epidural poly(pro-drug) thin films as a potential platform for sustained local pharmacotherapy after CNS injury.

