## Supplementary figures and images for "An Implantable Poly(pro-curcumin) Film for Local, Long-Lasting Neuroprotection After Spinal Cord Injury"

### Supp Figure 1

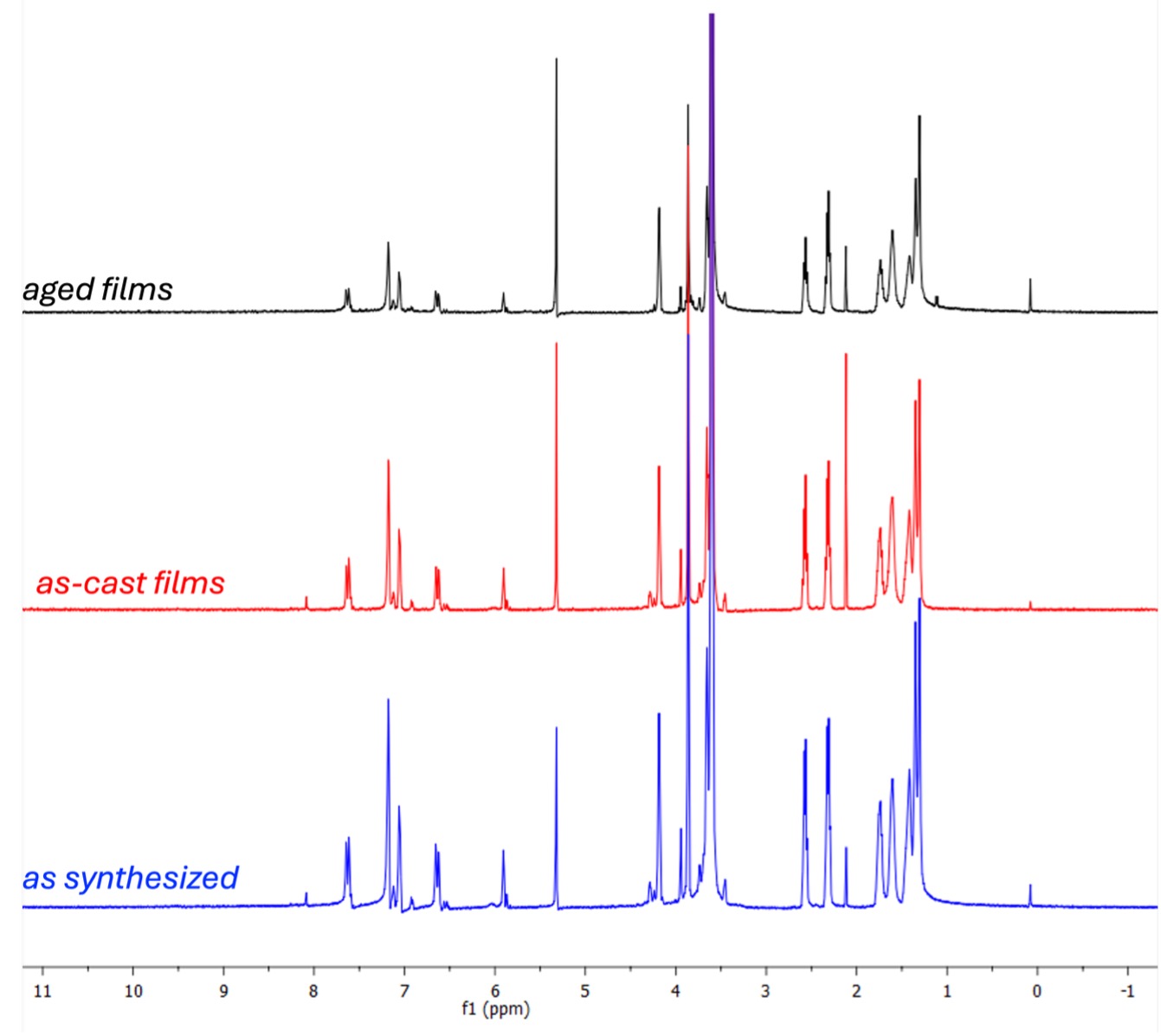

### Supp Figure 2

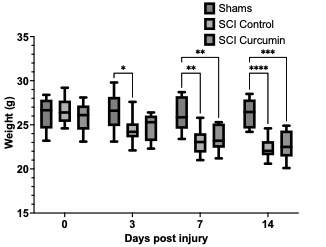

### Supp Figure 3

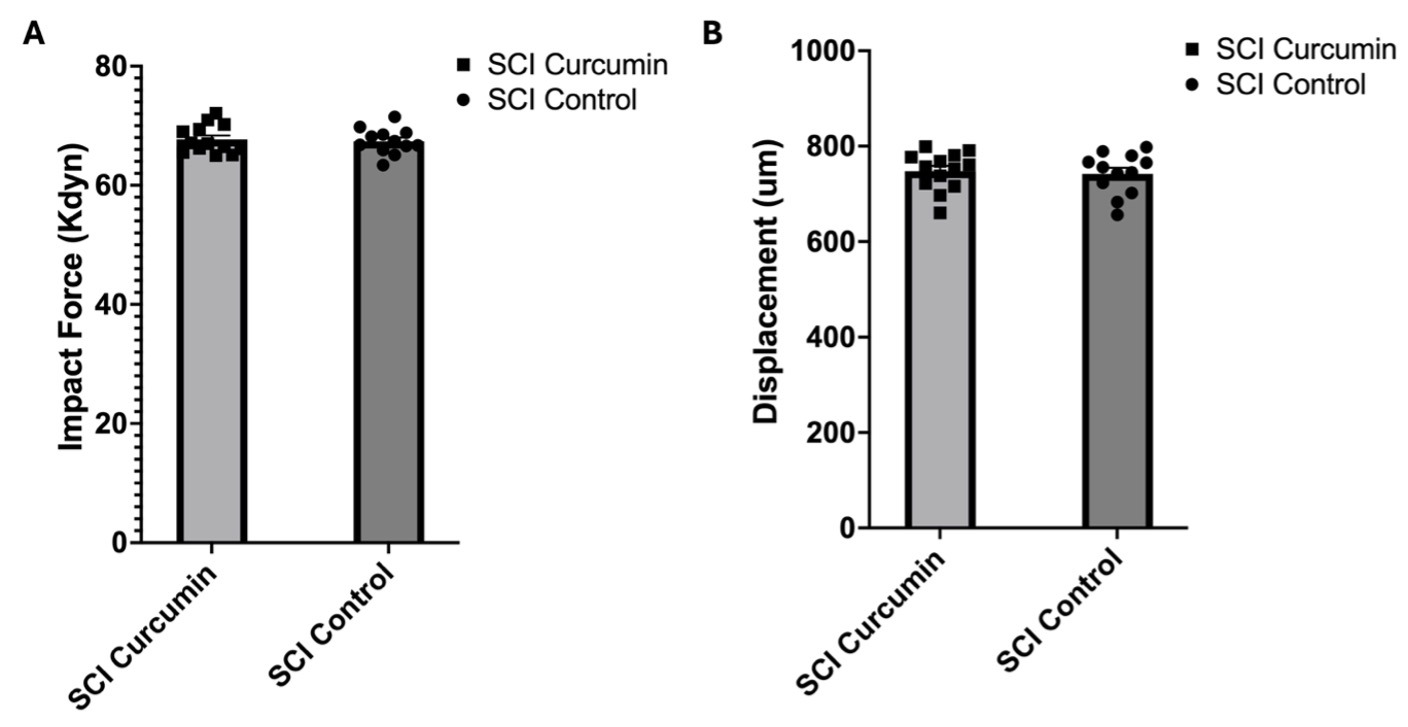
